# Learning dynamics of unsupervised deep learning reveal epoch-specific genetic architectures of brain morphology

**DOI:** 10.64898/2026.04.24.720631

**Authors:** Sheikh Muhammad Saiful Islam, Tian Xia, Xingzhong Zhao, Ziqian Xie, Degui Zhi

**Affiliations:** Department of Bioinformatics and Systems Medicine, McWilliams School of Biomedical Informatics (MSBMI) at University of Texas Health Science Center at Houston (UTHealth), USA

## Abstract

Representation learning is an emerging paradigm for deriving phenotypes from complex measurements (e.g., imaging) for genetic discovery. However, the learning dynamics of deep neural networks, especially the evolution of representations during training, while of interest in representation learning, were insufficiently investigated in the context of genetic discovery. In this study, using a 3D convolutional autoencoder trained on T1-weighted brain MRIs UK Biobank participants, we show that its learning trajectory forms an epoch-stratified landscape of brain morphology heritability. Different training epochs capture distinct genetic architectures at comparable heritability levels. Overall, ensembling across informative checkpoints identifies more genomic risk loci than the conventional single-checkpoint approach. Interpretability analysis reveals that epoch-specific loci, including MAPT and MCPH1, map onto biologically coherent and distinct neuroanatomical signatures, identified at different stages of the training process. Our results establish learning dynamics as a novel axis for genetic discovery using unsupervised deep learning and have practical implications for any architecture that saves multiple checkpoints during training.

## Introduction

Neuroimaging genetics aims to identify genetic variants that affect brain structure and function. It combines genome-wide association studies (GWAS) with brain imaging phenotypes. Large-scale efforts using UK Biobank and ENIGMA have identified hundreds of genomic loci associated with cortical and subcortical morphology. These efforts have improved our understanding of the brain’s genetic architecture (Adams et al., 2016; Elliott et al., 2018; Fan et al., 2022; Fu et al., 2024; García-Marín et al., 2024; Grasby et al., 2020; Hibar et al., 2015; Jansen et al., 2022; Makowski et al., 2023; Satizabal et al., 2019; Shadrin et al., 2021; S. M. Smith et al., 2021; Warrier et al., 2023). Traditionally, studies use imaging-derived phenotypes (IDPs) extracted from predefined brain parcellations. Software such as FreeSurfer and FSL is commonly used (Glasser et al., 2013; Miller et al., 2016; Perlaki et al., 2017). While useful, this approach is limited by the chosen atlas and segmentation pipeline. It can miss complex, distributed patterns of morphological variation that don’t fit predefined regions.

To overcome these limits, unsupervised deep learning models provide data-driven alternatives for extracting brain imaging phenotypes. This is especially true for autoencoders (Hinton & Salakhutdinov, 2006). These models compress entire MRI volumes into low-dimensional latent representations, without using anatomical priors. As a result, they can find features that reflect the full complexity of brain morphology. When used as endophenotypes for GWAS, such unsupervised representations have identified novel genomic loci not detected by traditional IDPs (Bonazzola et al., 2024; Patel et al., 2024), demonstrating their value for genetic discovery.

A key overlooked question is how an autoencoder’s genetic signals evolve during training. Standard practice selects a single checkpoint, usually the one with the lowest validation loss, while discarding other representations, assuming that superior reconstruction quality means greater genetic sensitivity. This assumption is rarely tested. Neural networks are known to undergo systematic changes in their internal representations across training (Rahaman et al., 2019), but whether these dynamics affect genetic discovery is unknown. Different training phases might encode distinct aspects of heritable brain morphology, in which case discarding all but one checkpoint would leave genuine genetic signals on the table.

Here, we systematically characterize the learning dynamics of a 3D convolutional autoencoder for genetic discovery and introduce TRACE (TRaining-dynamics Analysis via Checkpoint Ensemble), an epoch-ensembling strategy that leverages the entire training trajectory. We trained the model on T1-weighted brain MRIs from the UK Biobank, saving checkpoints at each of 85 epochs, and performed GWAS on 128-dimensional embeddings from an independent cohort of 22,962 individuals. The contributions are threefold: (1) we show that the learning trajectory forms a temporally structured landscape in which different epochs capture distinct genetic architectures of brain morphology at similar heritability, rendering the trajectory a multi-dimensional resource; (2) we develop a four-stage locus attribution pipeline—comprising epoch classification, integrated gradients, voxel-wise statistical mapping, and regional enrichment—that links epoch-specific genomic signals to anatomically interpretable brain regions; and (3) we demonstrate that epoch ensembling across informative checkpoints yields broader locus discovery than the conventional single-checkpoint approach at identical heritability. These findings demonstrate that the learning dynamics of unsupervised models provide new insights into the genetic architecture of brain morphology.

## Results

### Learning dynamics reveal divergent trajectories of reconstruction, prediction, and heritability

We trained a 3D convolutional autoencoder with a 128-dimensional bottleneck (138.12 million parameters) on 6,130 T1-weighted brain MRIs from the UK Biobank. We saved checkpoints after each of 85 epochs (Fig. 1a–b). During training, we used three metrics to assess representations: reconstruction loss, brain volume prediction, and SNP heritability (Fig. 1c). Reconstruction loss dropped rapidly in the first 10 epochs, then declined gradually until epoch 85, indicating progressive refinement of voxel-level image detail (Fig. 2d). Figure 2a–c shows example reconstructions and absolute difference maps for two subjects at epochs 3 (early), 37 (mid), and 83 (late). Even at epoch 3, despite high reconstruction loss, the autoencoder preserves global features such as cortical topology, ventricular geometry, and hemispheric structure (Fig. 2a–c). The absolute difference maps demonstrate that reconstruction errors remain diffuse and small throughout training. Across all epochs, the two subjects remain visually distinct, and morphological differences between subjects exceed reconstruction errors within each subject (Fig. 2a–c), confirming that embeddings preserve individual morphology from the earliest epochs.

**Figure 1.**
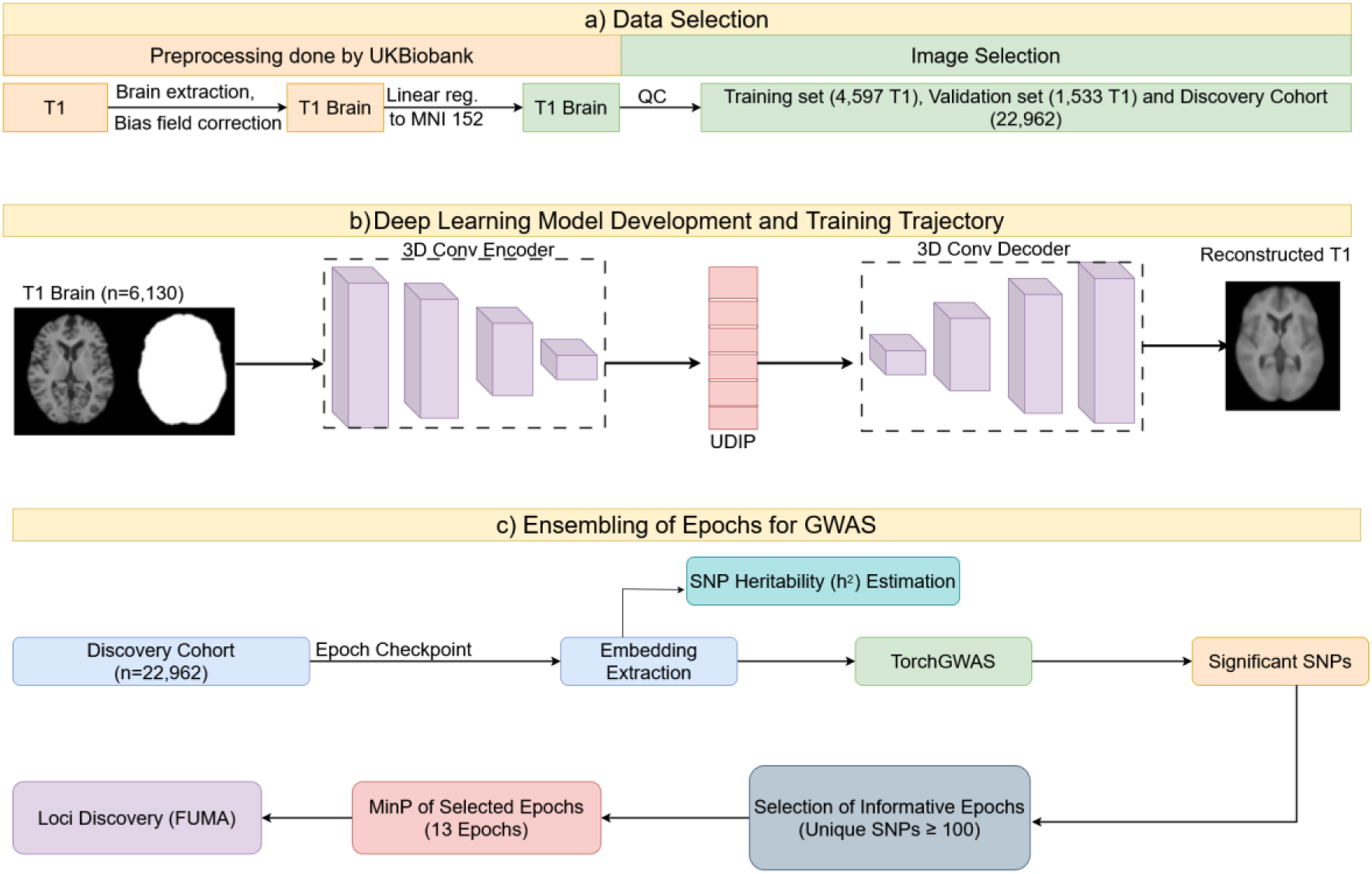
The TRACE framework pipeline. (a) Data selection: preprocessing of T1-weighted MRIs and cohort partitioning into training (n=6,130), validation (n=1,533) and independent discovery (n=22,962) sets. (b) Deep learning model development: 3D convolutional autoencoder architecture utilizing a whole-brain mask for MSE loss to generate unsupervised deep imaging phenotypes. (c) Epoch ensembling for GWAS: extraction of 128-dimensional embeddings across 85 epochs, followed by SNP heritability (h^2^) and selection of 13 informative checkpoints based on unique SNP discovery and final minP aggregation.

**Figure 2.**
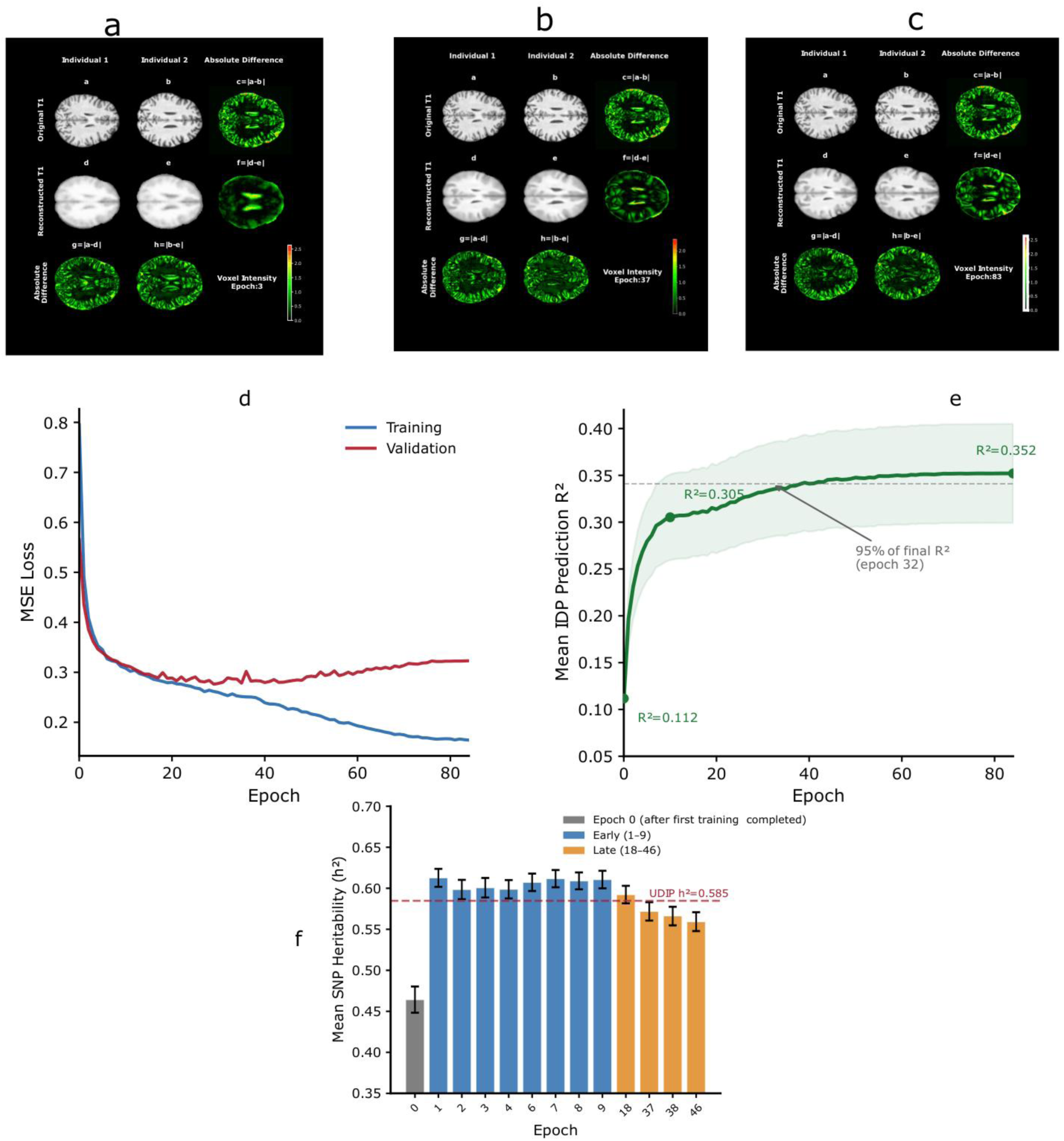
Training dynamics of the 3D convolutional autoencoder. (a–c) Representative MRI reconstructions for two subjects at epoch 3 (a, early training), epoch 37 (b, mid-training), and epoch 83 (c, near-convergence). Each panel shows the original MRI, reconstruction, and voxel-wise absolute difference map for both subjects. (d) Training and validation reconstruction loss (MSE) across 85 epochs. (e) Mean IDP prediction R² across 139 regional grey matter volumes as a function of epoch; shaded region = 95% CI across regions. (f) Mean SNP heritability (h²) at the 13 informative epochs, grouped by training stage: epoch 0 (dark blue), early epochs 1–9 (medium blue), and late epochs 18–46 (orange). Error bars = SD across 128 embedding dimensions.

To assess whether the learned embeddings capture anatomically meaningful information, we predicted 139 image-derived phenotypes (IDPs) that represent regional gray matter (GM) volumes. These GM volumes were measured independently by the UK Biobank imaging pipeline. We used 5-fold cross-validated ordinary least squares (OLS) regression on the 128-dimensional embeddings (n = 2,000 subjects). Even at epoch 0—after one complete pass through the training data—mean prediction R² across regions was 0.112. This suggests that, from the start, the learned representation from the convolutional architecture captures some spatial structure of brain morphology. By epoch 84, the mean prediction R² across regions reached 0.352. Mean R² rose steeply to 0.305 by epoch 10, representing 80% of the total improvement in just 12% of the training. It then reached 0.341 by epoch 40 and plateaued at 0.352 by epoch 84 (Fig. 2e; Supplementary Fig. 2; Supplementary Table 5). The best-predicted region was the Intracalcarine Cortex (R² = 0.554). This shows that 128 unsupervised dimensions can explain over half the variance in a specific regional brain volume. The improvement in IDP prediction was not uniform across regions. At late-training epochs (37, 38, 46; subsequently identified as MAPT-sensitive; see below), temporal lobe volumes showed a 69% relative increase in R² compared to early epochs (mean R² 0.347 vs 0.206), versus a 59% gain in non-temporal regions (0.340 vs 0.214; Mann-Whitney U P =0.031, Supplementary Fig. 5). The absolute difference in gain is modest (Δ 0.141 vs 0.126), and the grouping is coarse, so this analysis should be read as suggestive rather than definite. Nonetheless, the directional consistency between regions that show disproportionate IDP prediction gain and regions implicated in MAPT genetic signal is compatible with the interpretation that late-epoch representations preferentially encode temporal lobe morphology.

For heritability analyses on epochs that contributed independent genetic information, we first identified 13 informative epochs (0, 1, 2, 3, 4, 6, 7, 8, 9, 18, 37, 38, 46) using a data-driven criterion (see Methods: Epoch selection and ensembling for full details). SNP heritability (h²)—the proportion of phenotypic variance attributable to common genetic variants (Yang et al., 2011)—was then estimated for each of the 128 embedding dimensions at these 13 epochs using GCTA (Fig. 2f; Supplementary Fig. 1; Supplementary Table 1). Heritability was substantial at all epochs: mean h² = 0.464 at epoch 0 (after one complete pass through the training data), rising to h² ≈ 0.60–0.61 at epochs 1–9 (the highest values), and declining gradually to h² ≈ 0.56–0.57 at later epochs (37–46). All epoch-dimension pairs were highly significant (P < 1 × 10^−11^; Supplementary Table 1). The overall mean h² across all 13 epochs and 128 dimensions was 0.585 (SD = 0.136, n = 1,663 estimates), virtually identical to the h² of the single best validation epoch used in the conventional approach (0.585, SD = 0.134; see below).

These three metrics exhibit distinct trajectories during training. Reconstruction loss continues to improve, IDP prediction performance plateaus around epoch 40, and heritability peaks early, then slightly declines. This divergence is a crucial observation—demonstrating that the model’s ability to capture heritable variation does not align with its fidelity in reconstruction. Rather, the autoencoder’s learning dynamics establish a temporally structured landscape where genetic sensitivity evolves independently of image quality. Within this landscape, the regional specificity of IDP prediction gain corresponds to the regional specificity of epoch-specific genetic discovery; epochs that acquire MAPT sensitivity also demonstrate disproportionate improvement in encoding temporal lobe structures affected by MAPT.

#### The training trajectory reveals a temporally structured landscape of genetic discovery

After showing that reconstruction quality and heritability do not track each other across training, we next asked whether genetic signals captured also shift across training. We performed GWAS on 128-dimensional embeddings at each of the 85 training epochs using TorchGWAS(Zhao et al., 2026), a GPU-accelerated linear regression pipeline. This pipeline was applied to the discovery cohort (n = 22,962 White British individuals). For each epoch, we applied minimum P-value (minP) aggregation across the 128 dimensions to obtain a single genome-wide significance profile, using a Bonferroni-adjusted threshold of 5 × 10^−8^/128 per dimension.

The number of genome-wide significant SNPs varied substantially across epochs. While most epochs contributed significant associations, certain epochs yielded disproportionately more unique discoveries, defined as SNPs not detected at any earlier epoch. Applying a criterion of at least 100 unique SNPs per epoch, we identified 13 informative epochs: 0, 1, 2, 3, 4, 6, 7, 8, 9, 18, 37, 38, and 46 (Fig. 3c). These informative epochs collectively span the entire training trajectory, encompassing the initial post-optimization state (epoch 0) through early learning (epochs 1–9), mid-training (epoch 18), and late training (epochs 37–46).

**Figure 3.**
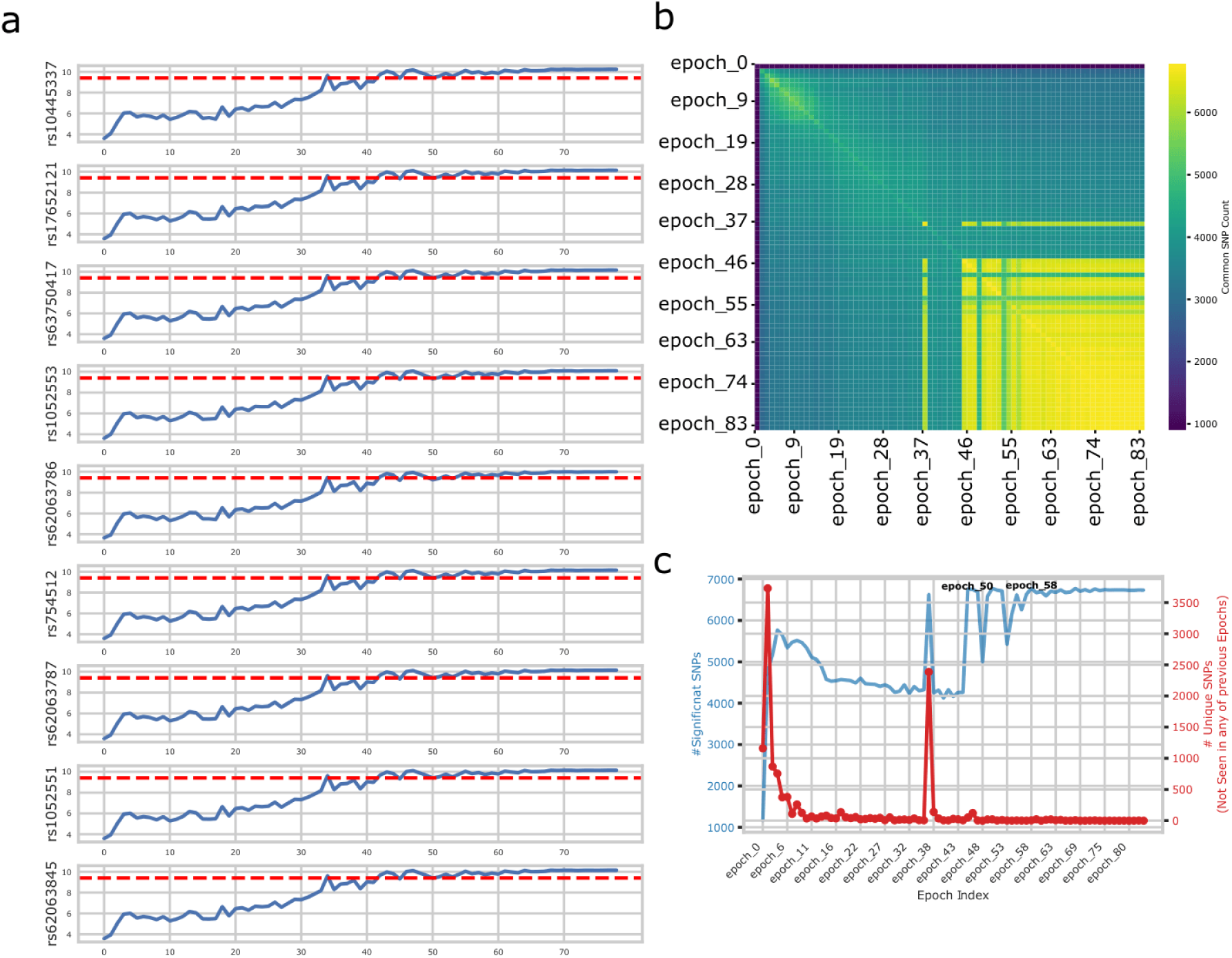
Epoch-specific genetic architectures. (a) Epoch-wise –log₁₀P values for select MAPT SNPs, showing detection only at later epochs. (b) Epoch-pair heatmap of shared significant SNPs. (c) Scatter plot of total significant SNPs versus unique SNPs per epoch, with 13 informative epochs annotated.

The epoch-pair heatmap of shared significant SNPs revealed that early epochs (0–9) share a largely overlapping set of associations, consistent with their similar heritability profiles. In contrast, later epochs (37 and beyond) exhibit a distinct set of associations that partially diverge from early-epoch findings (Fig. 3b). The later-epoch signal is primarily driven by the chr17q21.31 locus (lead SNP rs62062271, P = 9.75 × 10^−12^), a region of approximately 1.4 Mb encompassing MAPT and several co-localized genes, including KANSL1 and CRHR1, with the lead SNP falling within the MAPT gene body, a well-established locus associated with tau pathology and neurodegeneration. This locus is notably absent from early-epoch GWAS despite their higher overall heritability (h² ≈ 0.61 vs 0.57; Fig. 3a, Fig. 2f).

This divergence between heritability and locus discovery reflects the autoencoder’s learning dynamics across training. Early training captures coarse, globally distributed morphological variation, resulting in high total heritability but missing specific loci. As training progresses, the model refines its representations to capture finer morphological features, explaining the emergence of sensitivity to loci such as MAPT despite a slight reduction in total heritability. Thus, two representations with identical h² can encode different genetic signals, and the learning trajectory systematically shifts which genetic architectures are captured. This process produces a temporally structured landscape of complementary discoveries.

### Biological validation of epoch-specific sensitivity: the MAPT locus

To test if epoch-specific loci are genuine biological signals or just statistical artifacts, we performed anatomical mapping of the MAPT locus. The MAPT locus was the strongest example of epoch-specific detection. MAPT encodes the tau protein, which shows pathological aggregation in a clear neuroanatomical sequence. Neurofibrillary tangles start in the entorhinal cortex and extend from the temporal cortex to the parietal and frontal association areas (Vogel et al., 2020). If the autoencoder accurately captures MAPT-related brain morphology at later epochs, the relevant voxels should localize to temporal lobe structures.

We developed a four-stage locus mapping pipeline to answer a key question. Which brain regions does the autoencoder use when its representation becomes sensitive to the MAPT locus? First, to isolate the representational shift associated with MAPT detection, we trained an L1-penalized logistic regression classifier. This classifier distinguished embeddings from MAPT-negative epochs (0–36) from those of MAPT-positive epochs (60–83) using 500 randomly selected subjects per epoch. This process identified directions in embedding space that carry MAPT-relevant information.

Second, to generate voxel-level maps indicating which brain locations drive the classifier’s MAPT-positive prediction, we computed Integrated Gradient (IG) attribution maps through the full pipeline (MRI to encoder to embedding to classifier) for 30 subjects at each of 24 MAPT-positive epoch checkpoints. Results were averaged across epochs to get one saliency map per subject.

Third, to identify voxels consistently salient across subjects rather than those driven by individual variation, we performed a voxel-wise one-sample t-test across 30 subjects, applying FDR correction (α = 0.05). The resulting group-level saliency map revealed broad cortical involvement, with especially strong signal in inferior and lateral temporal regions at inferotemporal slices (Fig. 4a).

**Figure 4.**
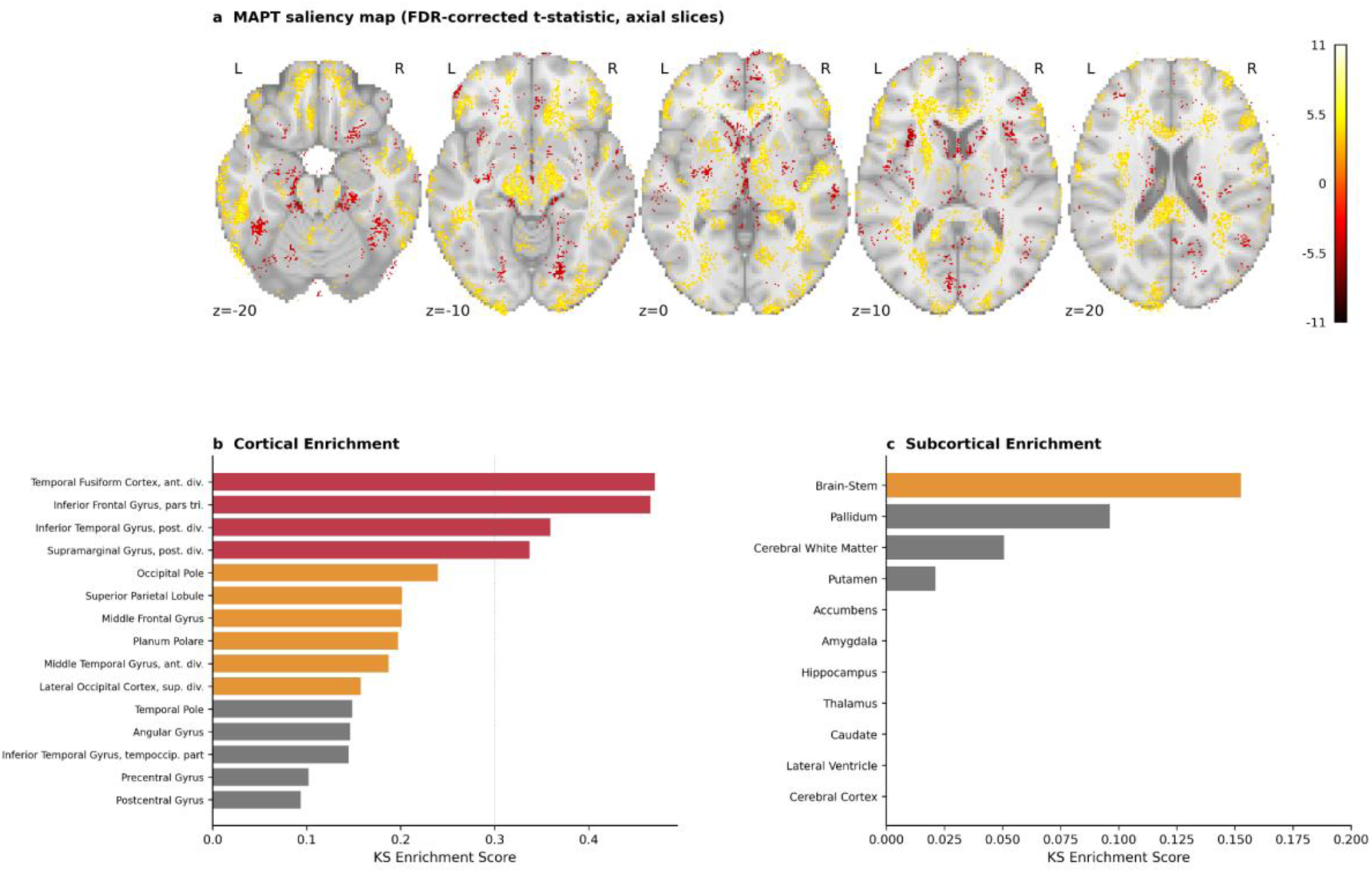
**MAPT locus biological validation**. The four-stage pipeline (epoch classification → Integrated Gradients → voxel-wise t-test → KS enrichment) is described in Methods. (a) Group-level FDR-corrected t-statistics saliency map (n=30 subjects) overlaid with MNI152 template, shown at five axial slices (z = −20, −10, 0, +10, +20 mm). Warm colors indicate voxels with consistently positive saliency across subjects. (b) KS enrichment scores for 47 Harvard–Oxford cortical regions, coloured by lobe; temporal lobe regions highlighted. (c) KS enrichment scores for 11 subcortical structures. Complete enrichment scores for all regions are in Supplementary Table 4.

Fourth, to test whether the saliency pattern is anatomically coherent and interpretable, we used Kolmogorov–Smirnov (KS) enrichment analysis with the Harvard–Oxford cortical and subcortical atlases (Jenkinson et al., 2012).

The enrichment analysis showed clear temporal lobe dominance (Fig. 4b; Supplementary Table 4). The highest-ranked cortical regions were the Temporal Fusiform Cortex, anterior division (KS = 0.471); Inferior Frontal Gyrus, pars triangularis (KS = 0.466); Inferior Temporal Gyrus, posterior division (KS = 0.359); and Supramarginal Gyrus, posterior division (KS = 0.337). Six of the top 13 enriched cortical regions were temporal lobe structures, including the Middle Temporal Gyrus (KS = 0.188), Planum Polare (KS = 0.198), and Temporal Pole (KS = 0.149). Subcortical enrichment was weak overall (Fig. 4c), with the highest KS score in the brainstem (0.153), consistent with tau pathology in progressive supranuclear palsy. In contrast, hippocampal enrichment was negligible (KS ≈ 0), which, although potentially counterintuitive given the hippocampus’s role in Alzheimer’s disease, aligns with large-scale GWAS evidence: the 17q21.31 locus affects cortical surface area (Grasby et al., 2020) but is not strongly linked with hippocampal volume in healthy adults (Hibar et al., 2015). This suggests that MAPT common variant effects on brain morphology are primarily cortical rather than hippocampal.

This anatomical mapping provides three layers of validation. First, the autoencoder learns biologically meaningful representations: voxel-level attribution through an unsupervised model recovers anatomically specific patterns that align with known tau neuroanatomy, despite the absence of genetic or anatomical labels during training. Second, epoch-specific loci represent genuine biological signals rather than noise; if late-epoch MAPT associations were statistical artifacts, the saliency maps would not exhibit coherent temporal lobe enrichment. Third, epoch ensembling captures biological signals that single checkpoints may miss. The MAPT locus illustrates the broader principle that different training epochs capture different loci, and no single checkpoint, regardless of selection criteria, can recover the entire genetic landscape. This finding supports a multi-epoch strategy that aggregates complementary signals from across the training trajectory. Notably, the spatial attribution maps reflect the full shift in representation between early and late training epochs. The enrichment of temporal lobe regions in these maps constitutes convergent evidence: two independent lines of evidence, genetic (MAPT detection at later epochs) and anatomical (late-epoch IG maps localizing to temporal lobe structures consistent with known MAPT neuroanatomy), converge on the same epochs.

### Biological validation of epoch-specific sensitivity: the MCPH1 locus

The MAPT locus is not the only example of epoch-specific biological sensitivity; it contrasts with the MCPH1 locus, which shows the opposite pattern during training. The MCPH1 locus (chr8p23.1, rs2979671, P = 2.56 × 10^−14^ at epoch 7) codes for microcephalin, a key protein that controls division of brain precursor cells. Mutations here cause a genetic brain disorder characterized by a much smaller cortical surface area and highly abnormal brain folding (Passemard et al., 2013). While MAPT appears only at later epochs (37+), MCPH1 is significant only during early training (epochs 2–12) and is not detected thereafter. These two loci show a sharp contrast: distinct genetic signals are observed at opposite ends of the training period.

We used the same four-stage pipeline for the MCPH1 locus that we used for the MAPT locus, where the MCPH1-positive encodings are from epochs 2-12, while the MCPH1-negative embeddings are from epochs 37-83. A voxel-wise one-sample t-test with FDR correction (α = 0.05) produced the group-level saliency map (Supplementary Figure 3a), and KS enrichment against Harvard-Oxford atlases provided regional interpretation.

The cortical enrichment pattern differed markedly from MAPT, showing no temporal lobe dominance (see Supplementary Figure 3b; Supplementary Table 6). In contrast, enrichment was concentrated in a perisylvian cluster, including the Supracalcarine Cortex (KS = 0.119), Parietal Operculum Cortex (KS = 0.107), Parahippocampal Gyrus, posterior division (KS = 0.090), Paracingulate Gyrus (KS = 0.089), Frontal Operculum Cortex (KS = 0.084), Insular Cortex (KS = 0.083), Central Opercular Cortex (KS = 0.076), and Heschl’s Gyrus (KS = 0.069). Subcortical regions, such as the Caudate (KS = 0.064) and Amygdala (KS = 0.056), also showed strong enrichment (see Supplementary Figure 3c). Notably, this perisylvian pattern is biologically grounded: development around the Sylvian fissure and insula is especially disrupted in MCPH1-related primary microcephaly (Passemard et al., 2013). Consistent with these findings, the MCPH1 common variant has also been reported to associate with whole-brain MRI metrics (Fan et al., 2022).

The spatial dissociation between MCPH1 (perisylvian/opercular) and MAPT (temporal lobe) enrichment patterns underscores a key contrast: epoch-specific genetic signals correspond to distinct morphological processes. Notably, two independent loci, detected at opposite ends of the training trajectory, display non-overlapping anatomical enrichment profiles, each of which aligns with the established biology of its respective gene. This convergence strengthens the claim that TRACE epoch specificity reflects genuine biological variation, clarifying that MAPT’s temporal lobe enrichment stems from the locus’s neuroanatomy, rather than representing a general property of late-epoch features.

### Harnessing learning dynamics through epoch ensembling

TRACE constructs its ensemble by first selecting epochs that each contribute at least 100 genome-wide significant SNPs not detected at any earlier epoch—a data-driven criterion that identified 13 informative epochs (0, 1, 2, 3, 4, 6, 7, 8, 9, 18, 37, 38, 46; Fig. 3c)—and then aggregating their GWAS summary statistics across all 128 embedding dimensions via minP, retaining the smallest p-value observed for each SNP across the full set of 1,664 epoch–dimension combinations. To quantify the advantage of this approach, we compared TRACE against UDIP ((Patel et al., 2024)), an established single-checkpoint method that uses the same autoencoder architecture but selects only the best validation epoch (lowest reconstruction loss) for GWAS—the standard practice in deep learning-based neuroimaging genetics. Both methods use minP aggregation across 128 embedding dimensions, and loci were annotated using FUMA. Epoch ensembling identified 90 genomic risk loci (Supplementary Table 2), compared with 78 from UDIP (Supplementary Table 3). Using interval-based overlap on the same chromosome, 54 loci were shared between the two methods, 36 were unique to the epoch ensemble, and 24 were unique to UDIP (Fig. 5b). Critically, neither approach strictly dominates the other: the two methods access overlapping but distinct regions of the genetic landscape, reflecting their different selection criteria. All 36 ensemble-unique loci reached genome-wide significance with strong P-values (many P < 4.88 × 10^−14^), confirming they are genuine associations rather than artifacts of the more conservative ensemble threshold.

**Figure 5.**
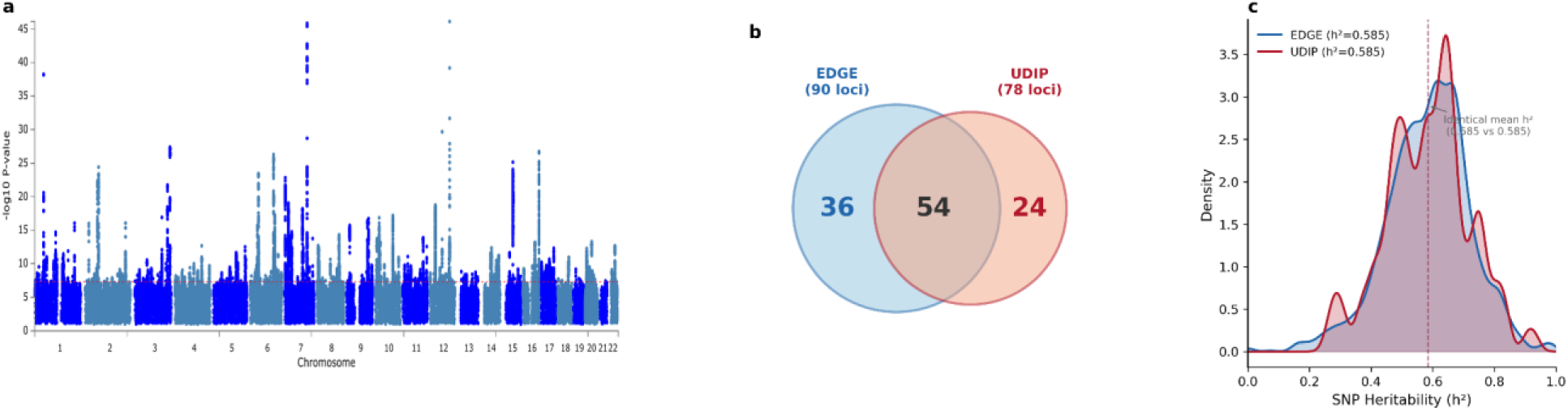
TRACE vs UDIP comparison. (a) Manhattan plot of the ensemble TRACE GWAS, showing 90 genomic risk loci. (b) Venn diagram: 54 shared, 36 TRACE-unique, 24 UDIP-unique loci. (c) Heritability density plot: TRACE and UDIP h² distributions overlaid, showing near-identical means.

To assess the robustness of the ≥100 unique SNP criterion, we repeated the analysis using thresholds of 50 and 200 unique SNPs. When we applied a threshold of 50 unique SNPs, we obtained 19 informative epochs at an adjusted Bonferroni threshold of 2.06 × 10^−11^. Using a higher threshold of 200 unique SNPs resulted in 8 informative epochs, with a Bonferroni threshold of 4.88 × 10^−11^. These analyses discovered 86 and 90 genomic loci, respectively. In comparison, the primary threshold identified 90 loci. Together, these results demonstrate that the main conclusions are robust across alternative epoch selection criteria (Supplementary Figure 4; Supplementary Table 7).

Critically, the mean SNP heritability was virtually identical between the two approaches: UDIP h² = 0.585 (SD = 0.134, 128 dimensions) versus TRACE grand mean h² = 0.585 (SD = 0.136, 1,663 dimension-epoch pairs; Fig. 5c). This demonstrates that the advantage of epoch ensembling is not higher heritability but broader genetic coverage: different checkpoints capture different heritable architectures of brain morphology at comparable levels of total genetic variance explained.

The 24 UDIP-unique loci—present in UDIP but absent from the epoch ensemble—deserve explicit attention, as they reveal the trade-off inherent in epoch selection. The 13 informative epochs were selected by a data-driven criterion (≥100 unique SNPs per epoch), which favors epochs that introduce many novel associations relative to all prior epochs. The validation-loss-optimal UDIP epoch is selected by a fundamentally different criterion—reconstruction quality—and carries its own genetic profile not necessarily captured by any of the 13 informative epochs. These are not missing signals but signals with a different discovery path: the UDIP epoch maximizes reconstruction fidelity, which may preferentially encode certain morphological features whose genetic architecture is not among the most temporally novel. Epoch ensembling and single-checkpoint approaches therefore have complementary biases, and the full landscape of brain morphology heritability likely requires both strategies. Denser epoch sampling or adaptive selection algorithms that explicitly optimize for UDIP-complementary coverage could reduce this gap in future work.

The MAPT locus (chr17q21.31) was detected by both methods through overlapping genomic intervals, but the epoch-resolved analysis provides additional insight: MAPT is only detectable at later epochs (37+), not at early epochs. This means UDIP detects MAPT because its best validation epoch is sufficiently late, whereas epoch ensembling explicitly reveals the epoch-specificity of this detection. Early-epoch GWAS with higher h² (≈0.61) fail to detect MAPT, suggesting that heritability and locus discovery can diverge.

To validate that the discovered loci reflect genuine genetic signals, we tested replication of all 114 loci (90 TRACE + 24 UDIP-unique) in an independent cohort of 12,258 unrelated UK Biobank participants (kinship-pruned across discovery and replication). For each locus, the lead SNP from FUMA was tested at its discovery-best epoch — the epoch at which the association was strongest in discovery — in the replication TorchGWAS. Replication was defined as p < 0.05 with a consistent effect direction. All 36 TRACE-unique, all 54 shared, and all 24 UDIP-unique loci replicated nominally (114/114; 100%), with 100% direction concordance. Applying a Bonferroni correction for loci tested (0.05/114 = 4.4 × 10^−4^), 24/36 TRACE-unique loci (66.7%), 53/54 shared loci (98.1%), and 14/24 UDIP-unique loci (58.3%) replicated (Supplementary Figure 7;Supplementary Table 8). The higher Bonferroni rate for shared loci reflects their detection by two independent methods; the lower rates for method-specific loci are expected, given the replication sample is approximately half the discovery size. Epoch-specificity was also reproducible: lead SNPs for MAPT and MCPH1 were nominally significant at 2/2 and 7/7 of their respective discovery-informative epochs in the replication cohort, with 100% effect direction concordance at all epochs (Supplementary Table 9; Supplementary Figure 6).

## Discussion

This study demonstrates that the learning dynamics of an unsupervised autoencoder generate a temporally structured landscape of heritable variation in brain morphology. As training progresses, the model captures different sets of loci at each stage, with limited overlap between early and later epochs. This means the learning trajectory explores complementary regions of the genetic landscape, rather than repeatedly detecting the same signals. The central finding is that this dynamic can be exploited. Epoch ensembling across 13 informative checkpoints identifies 90 genomic loci, while the conventional single-checkpoint approach yields only 78. Notably, 36 loci are unique to the ensemble and 24 to the single checkpoint, both at identical mean heritability (h² = 0.585). The two approaches thus exhibit complementary strengths—each captures signals missed by the other. The ensemble, however, accesses a broader region of the genetic landscape. The fundamental advantage of ensembling lies in broader, more diverse genetic coverage, not in increased heritability; this is a direct result of the learning dynamics.

These findings challenge the implicit assumption that the best-reconstructing checkpoint is optimal for genetic analysis. Specifically, reconstruction loss, anatomical prediction accuracy, and SNP heritability do not track one another across training: reconstruction loss improves continuously, IDP prediction plateaus at approximately epoch 40, heritability peaks early and then declines modestly, and genomic loci continue to emerge at later epochs even as the other two metrics stabilize. This pattern is consistent with the progressive feature refinement observed broadly in deep learning (Rahaman et al., 2019), in which networks tend to first capture coarse, global structure before resolving finer details — a transition our results suggest has direct consequences for genetic discovery. Whether epoch-specific genetic architectures reflect frequency-domain specialization or other aspects of the loss landscape geometry remains an open question.

The MAPT locus mapping provides biological validation of genetic sensitivity at specific training epochs through two independent lines of evidence. First, the MAPT genetic signal appears only at late training epochs. Second, when we compute attribution maps (Integrated Gradients distinguishing early- from late-epoch embeddings across representational dimensions), the saliency localizes to the temporal lobe, the site of tau pathology. The classifier was trained on overall representational change, not MAPT-specific effects, so temporal lobe enrichment in these maps is an independent anatomical signal. The dominance of the temporal fusiform cortex and inferior temporal gyrus matches the pattern of tau pathology described by Braak staging (Vogel et al., 2020). The simultaneous emphasis on the inferior frontal gyrus reflects the frontotemporal pattern seen in frontotemporal dementia (Rascovsky et al., 2011; Strang et al., 2019). The convergence of these two independent forms of evidence — genetic and anatomical — at the same epochs shows that epoch-specific loci are real biological signals, not artifacts. This pattern extends to another locus at the start of training: MCPH1 (chr8p23.1), detected only at early epochs (2–12), with attribution maps localized to a perisylvian/opercular cluster, in line with microcephalin’s known role in perisylvian brain folding. Therefore, MAPT does not have undue evidential weight for the general claim.

Checkpoint ensembling has been explored in cardiac MRI phenotyping (Bonazzola et al., 2024). However, the learning dynamics perspective we present here—where we systematically characterize how genetic sensitivity evolves across training epochs—is, to our knowledge, novel in neuroimaging genetics. The complexity of the brain’s structural variability (differences in brain shapes or structures across individuals) and its polygenic genetic architecture make this domain well-suited to reveal epoch-dependent sensitivity at different training times. Our approach complements existing multivariate GWAS methods, such as MOSTest (van der Meer et al., 2020). Those methods aggregate across phenotypic dimensions (combining multiple traits), while we aggregate across training stages (combining results from different points in model training).

The practical implications extend beyond our specific autoencoder. Any unsupervised deep learning model used for genetic phenotyping undergoes learning dynamics that may create epoch-specific sensitivity to different genetic architectures. Our results suggest it is useful to systematically characterize these dynamics, rather than relying solely on validation loss for checkpoint selection. This strategy could improve genetic discovery across a range of architectures, such as variational autoencoders (Kingma et al., 2019) and transformers (Vaswani et al., 2017), and across imaging modalities, including diffusion MRI and functional MRI. The framework is architecture-agnostic and only requires that multiple checkpoints be saved during training.

Several limitations should be noted. First, our analysis used 13 of 85 available epochs, selected by a data-driven criterion (≥100 unique SNPs). Sensitivity analyses with thresholds of 50 and 200 unique SNPs yielded 86 and 90 loci, respectively, confirming robustness of the primary result. The 24 UDIP-unique loci reflect the complementary bias of reconstruction-quality-based checkpoint selection. Combining selection criteria or denser epoch sampling could improve joint coverage. Second, IG attribution at 24 checkpoints x 30 subjects was already computationally expensive; scaling to hundreds of subjects per checkpoint would require roughly ∼1,000 GPU-hours and remains the most straightforward path to tighter spatial maps. Third, the KS enrichment analysis is descriptive and does not provide formal significance testing for individual regions. Future work could employ permutation-based enrichment. Fourth, our study uses only T1-weighted structural MRI from the UK Biobank’s White British cohort. Generalizability to other imaging modalities, ancestries, and cohorts will require external validation. Fifth, IDP prediction used OLS regression. Nonlinear methods might reveal more anatomical information in the embeddings.

In summary, the learning dynamics of unsupervised deep learning models provide a window into the genetic architecture of brain morphology. The training trajectory forms a temporally structured landscape: different epochs capture distinct heritable variation, and no single checkpoint fully represents this landscape. Epoch ensembling thus offers a principled way to harness these dynamics, achieving broader genetic coverage at identical heritability. Together, these findings establish learning dynamics as a new axis of investigation in deep learning-based neuroimaging genetics.

## Methods

### Dataset

We used T1-weighted brain MRI data from the UK Biobank (Alfaro-Almagro et al., 2018; Littlejohns et al., 2020; Sudlow et al., 2015). MRIs were acquired primarily on Siemens Skyra 3T scanners with a 32-channel RF head coil at 1 × 1 × 1 mm resolution. Preprocessing followed the UK Biobank imaging pipeline: defacing, brain extraction (FSL BET), bias field correction (FSL FAST), and affine registration (12 degrees of freedom) to MNI152 standard space using FSL FLIRT. Intensity values were z-score normalized, excluding background voxels.

Quality control used UK Biobank metrics: inverted contrast-to-noise ratio (field 25735) and T2/T1 discrepancy (field 25736). Images within the top 95% of quality were retained. Only one visit per participant was included, yielding 6,130 images randomly split into training (n = 4,597, 75%) and validation (n = 1,533, 25%) sets.

An independent cohort of 35,810 White British participants (UK Biobank fields 21000 and 22006) with T1-weighted MRIs—not part of the deep learning training—was used for GWAS. These were divided into a discovery cohort (n = 22,962) and a replication cohort (n = 12,848).

### 3D convolutional autoencoder

We employed a 3D convolutional autoencoder to learn 128-dimensional representations of endophenotypes from whole-brain T1-weighted MRIs. The architecture consists of an initial convolutional block, four encoder blocks, a 128-dimensional fully connected latent space, four decoder blocks, and a final convolutional block, totaling 138.12 million parameters.

Each encoder block comprises a 3D max-pooling layer (kernel size = 2) followed by two 3D convolutional layers (kernel size = 3, stride = 1), batch normalization, and LeakyReLU activation. The decoder mirrors the encoder using 3D transposed convolutions (stride = 2, kernel size = 2). Skip connections were intentionally omitted to ensure full information compression into the latent space. The model processes entire MRI volumes (182 × 218 × 182) at native resolution, allowing each latent dimension to encode whole-brain information.

### Training

The model was trained as a voxel-wise regression task minimizing mean squared error (MSE) between input and reconstructed images, with a brain mask excluding non-brain voxels from the loss computation. We used the AdamW optimizer with the OneCycleLR learning rate scheduler (L. N. Smith, 2018). Training was conducted for 85 epochs on a single NVIDIA A100-SXM-80GB GPU, with each epoch (training + validation) taking approximately 30 minutes. Checkpoints were saved at every epoch to enable systematic analysis of the training trajectory.

### Genome-wide association study

Genome-wide scans were conducted using approximately 10 million SNPs, including 658,720 directly genotyped SNPs (UK BiLEVE Axiom Array) and imputed variants. For each of the 85 training epochs, we extracted 128-dimensional embeddings from the discovery cohort and performed GWAS using TorchGWAS, a GPU-accelerated linear regression pipeline developed in our laboratory. Age, sex, and the first 10 genetic principal components were included as covariates. A genome-wide significance threshold of 5 × 10^−8^/128 (Bonferroni correction for 128 dimensions) was applied per epoch.

### Epoch selection and ensembling

To select informative epochs for ensembling, we identified epochs that contributed ≥100 genome-wide significant SNPs not detected at any earlier epoch. This data-driven criterion yielded 13 informative epochs: 0, 1, 2, 3, 4, 6, 7, 8, 9, 18, 37, 38, and 46. For each informative epoch, the minimum P-value (minP) across 128 dimensions was computed for every SNP, and results were aggregated across the 13 epochs using a second-level minP, with a Bonferroni-corrected significance threshold of 5 × 10^−8^/ (128 × 13) = 3.00 × 10^−11^, accounting for both embedding dimensions and selected epochs. This threshold is more conservative than the per-epoch threshold applied to UDIP (5 × 10^−8^/128 = 3.91 × 10^−10^), ensuring the ensemble comparison is conservative against false positives. Genomic risk loci were annotated using FUMA (Watanabe et al., 2017).

We used SNP counts rather than locus counts to retain sensitivity to association signals that have not yet been consolidated into annotated loci. We defined novelty sequentially relative to all prior epochs. This approach directly measures marginal information gain without requiring prior knowledge of the embedding geometry.

### SNP heritability estimation

SNP heritability (h²) was estimated for each of the 128 embedding dimensions at each of the 13 informative epochs using GCTA v1.94 (Jiang et al., 2019). We used the fastGWA mixed linear model with the --model-only flag, which estimates variance components (Vg, Ve) from a sparse genomic relationship matrix (GRM) without performing genome-wide SNP scanning. Heritability was computed as h² = Vg/(Vg + Ve). Age, sex, and the first 10 genetic principal components were included as covariates. This yielded 1,664 heritability estimates (13 epochs × 128 dimensions).

For comparison, UDIP heritability was obtained from the full fastGWA-mlm analysis of the best validation epoch, providing 128 h² estimates from the single-checkpoint approach.

### IDP prediction

We assessed the anatomical information encoded in the 128-dimensional embeddings by predicting 139 regional grey matter (GM) volumes from the UK Biobank imaging data (fields 25782–25920): 96 cortical, 15 subcortical, and 28 cerebellar volumes, derived from the Harvard–Oxford atlas. For each epoch (0–84) and each GM volume, we performed 5-fold cross-validated OLS regression (n = 2,000 subjects), standardizing embeddings and volumes within each training fold to prevent data leakage. Prediction accuracy was quantified as R² (squared Pearson correlation between predicted and actual values).

### MAPT locus mapping

#### Epoch classification

We trained an L1-penalized logistic regression classifier to distinguish MAPT-negative (epochs 0–36) from MAPT-positive (epochs 60–83) embeddings. For each labeled epoch, 500 subjects were randomly selected, yielding 30,500 training samples (61 epochs × 500 subjects). Features were standardized, and the classifier was fit with an 80/20 stratified train/test split.

#### Integrated Gradients attribution

For 30 randomly selected subjects, we computed voxel-level attribution maps using Integrated Gradients (Sundararajan et al., 2017) through the full prediction pipeline: T1 MRI → autoencoder encoder → 128-dim embedding → standard scaler → classifier logit. The baseline was the average of 30 ‘brainless’ images (with foreground voxels replaced by the background mean). Attribution maps were computed at each of 24 MAPT-positive epoch checkpoints (n_steps = 50) and averaged across epochs, yielding one saliency map per subject. We note that the classifier was trained to distinguish early- from late-epoch embeddings across all representational dimensions, not specifically on MAPT. Accordingly, the resulting attribution maps reflect the aggregate representational shift between early and late training epochs — the voxels that drive ‘late-epoch’ classification — rather than a direct attribution of MAPT genetic effects to specific voxels.

### Statistical analysis

Saliency maps were stacked into a 4D array (30 × 182 × 218 × 182) and subjected to a voxel-wise one-sample t-test (H₀: mean saliency = 0), with background voxels masked out. P-values were corrected using Benjamini–Hochberg FDR at α = 0.05.

### Regional enrichment

We quantified enrichment of the MAPT saliency signal in anatomical regions using a KS enrichment analysis analogous to Gene Set Enrichment Analysis (Subramanian et al., 2005). Voxels covered by the Harvard–Oxford cortical atlas (48 regions) and subcortical atlas (11 structures), thresholded at 50%, were ranked by t-statistic. For each region, a running enrichment score was computed by walking down the ranked list, incrementing when a voxel belongs to the target region and decrementing otherwise. The KS score is the maximum of this running sum. Left and right hemispheres were merged.

### Locus comparison

TRACE and UDIP genomic risk loci (from FUMA) were compared using interval-based overlap: two loci are considered “shared” if they are on the same chromosome and their genomic intervals [start, end] overlap (i.e., start₁ ≤ end₂ and start₂ ≤ end₁). Loci present in only one method were classified as “unique” to that method.

### Usage of LLM

Claude Code (Claude Sonnet 4.6, Anthropic) was used in the preparation of this manuscript. Its use was limited to converting author-produced numerical outputs and analytical conclusions into draft prose; all results, interpretations, and scientific conclusions were generated entirely by the authors using the computational pipelines described in this Methods section. All draft text produced by the tool was subsequently reviewed, substantially revised, and approved by all authors, who bear full responsibility for the accuracy and integrity of the final manuscript.

### Data availability

UK Biobank data are available through the UK Biobank Access Management System (https://www.ukbiobank.ac.uk). GWAS summary statistics and FUMA annotations will be made available at Zenodo upon publication.

### Code availability

Code for the TRACE pipeline, including the autoencoder architecture, TorchGWAS pipeline, epoch ensembling, and MAPT locus mapping, is in https://github.com/ZhiGroup/TRACE and will be made available on GitHub upon publication.

## Acknowledgements

This research has been conducted using the UK Biobank Resource under Application Number 24247. We thank the UK Biobank participants and the UK Biobank team for making this resource available.

## Author contributions

S.M.S.I., Z.X. and D.Z. conceived the study. S.M.S.I designed and implemented the deep learning pipeline, conducted all GWAS analyses, and performed all downstream computational analyses, with substantial input from D.Z, and Z.X. S.M.S.I., Z.X. and D.Z led the result interpretation, and D.Z. secured the funding for the project. D.Z. oversaw the execution of the entire project. S.M.S.I and D.Z. led the writing of the manuscript, with substantial inputs from Z.X., T.X. and X.Z. All authors discussed, commented and confirmed the final version of the manuscript.

## Ethics declarations

Our analysis was approved by the UTHealth committee for the protection of human subjects under No. HSC-SBMI-20-1323. UKBB has secured informed consent from the participants in the use of their data for approved research projects. UKBB data was accessed via approved project 24247.

## Supplementary Information

### Supplementary Figures

Supplementary Figure 1: Distribution of SNP heritability (h²) across 128 embedding dimensions for each of the 13 informative epochs, shown as violin plots. The dashed red line indicates the UDIP mean h² = 0.585.

Supplementary Figure 2: IDP prediction R² heatmap across training epochs for the top 20 best-predicted regional GM volumes, showing the progressive improvement in anatomical encoding from epoch 0 to epoch 84.

Supplementary Figure 3: MCPH1 locus biological validation. The same four-stage pipeline was applied to the MCPH1 locus (MCPH1-positive: epochs 2–12; MCPH1-negative: epochs 37–83). (a) Group-level FDR-corrected t-statistics saliency map (n=30 subjects) overlaid with MNI152 template. (b) KS enrichment scores for 47 Harvard–Oxford cortical regions, with perisylvian/opercular regions highlighted. (c) KS enrichment scores for 11 subcortical structures. Complete enrichment scores are in Supplementary Table 6.

Supplementary Figure 4: Sensitivity analysis of the epoch selection threshold. Genomic loci identified under three unique-SNP thresholds (≥50, ≥100 [primary], ≥200). Results are stable across thresholds (86, 90, and 90 loci, respectively).

Supplementary Figure 5: Epoch-specific convergence of genetic and anatomical sensitivity. (a) Mean IDP prediction R² trajectory for temporal (red) vs non-temporal (blue) GM volumes across 85 epochs. (b) Mean R² at early, late-training, and final epochs; temporal lobe gain is significantly greater (Mann–Whitney U P = 0.031).

Supplementary Figure 6: Epoch-specificity replication. Per-epoch -log₁₀(P) profiles in discovery (N ≈ 23k; left) and replication (right) for MAPT, MCPH1, and the five top-ranked TRACE-unique loci. Red bars indicate discovery-informative epochs (p < 3 × 10⁻¹¹). Red horizontal line = ensemble Bonferroni threshold. Spearman ρ and direction concordance are annotated on each replication panel.

Supplementary Figure 7: Replication of all 114 discovered loci in an independent cohort (N ≈ 12k). (a) Nominal and Bonferroni replication rates by locus category (TRACE-unique, Shared, UDIP-unique). All 114 loci replicated nominally (100%); shared loci replicated at the highest Bonferroni rate (98.1%), reflecting detection by two independent methods. (b) Scatter plot of discovery vs replication −log₁₀(P) for all 114 loci, coloured by category. Dashed lines indicate nominal (p = 0.05) and Bonferroni (p = 4.4 × 10⁻⁴) thresholds. A stronger discovery signal predicts a stronger replication signal, consistent with a power explanation for the lower Bonferroni rate among method-specific loci.

### Supplementary Tables

Supplementary Table 1: Full heritability results for all 13 informative epochs, including per-epoch mean h², standard deviation, median, number of valid dimensions, and mean standard error.

Supplementary Table 2: Complete list of 90 TRACE genomic risk loci with genomic coordinates, lead SNPs, P-values, and overlap status with UDIP loci.

Supplementary Table 3: Complete list of 78 UDIP genomic risk loci with genomic coordinates, lead SNPs, P-values, and overlap status with TRACE loci.

Supplementary Table 4: KS enrichment scores for all 47 cortical and 11 subcortical Harvard–Oxford atlas regions from the MAPT saliency analysis.

Supplementary Table 5: IDP prediction R² for all 139 regional grey matter volumes at selected epochs (0, 10, 40, 84), with anatomical category annotations.

Supplementary Table 6: KS enrichment scores for all 47 cortical and 11 subcortical Harvard–Oxford atlas regions from the MCPH1 saliency analysis, parallel to Supplementary Table 4.

Supplementary Table 7: Sensitivity analysis of the epoch selection threshold. Genomic loci (post-FUMA) were identified under three unique-SNP thresholds (≥50, ≥100, ≥200) with corresponding epoch counts, Bonferroni thresholds, and locus counts.

Supplementary Table 8: Per-locus replication results for all 114 loci (36 TRACE-unique, 54 shared, 24 UDIP-unique). Columns: category, locus coordinates, lead SNP, discovery-best epoch, discovery P, replication P, effect direction concordance, nominal replication status, Bonferroni replication status.

Supplementary Table 9: Epoch-specificity replication for MAPT, MCPH1, and five top-ranked TRACE-unique loci. Columns: locus, lead SNP, discovery-informative epochs, Spearman ρ (epoch profile concordance), most_sig_dim concordance at informative epochs, direction concordance, and nominal replication rate at informative epochs.

## Reference

Adams, H. H. H., Hibar, D. P., Chouraki, V., Stein, J. L., Nyquist, P. A., Rentería, M. E., Trompet, S., Arias-Vasquez, A., Seshadri, S., Desrivières, S., Beecham, A. H., Jahanshad, N., Wittfeld, K., Van der Lee, S. J., Abramovic, L., Alhusaini, S., Amin, N., Andersson, M., Arfanakis, K., … Thompson, P. M. (2016). Novel genetic loci underlying human intracranial volume identified through genome-wide association. Nature Neuroscience, 19(12), 1569–1582.

Alfaro-Almagro, F., Jenkinson, M., Bangerter, N. K., Andersson, J. L. R., Griffanti, L., Douaud, G., Sotiropoulos, S. N., Jbabdi, S., Hernandez-Fernandez, M., Vallee, E., Vidaurre, D., Webster, M., McCarthy, P., Rorden, C., Daducci, A., Alexander, D. C., Zhang, H., Dragonu, I., Matthews, P. M., … Smith, S. M. (2018). Image processing and Quality Control for the first 10,000 brain imaging datasets from UK Biobank. NeuroImage, 166, 400–424.

Bonazzola, R., Ferrante, E., Ravikumar, N., Xia, Y., Keavney, B., Plein, S., Syeda-Mahmood, T., & Frangi, A. F. (2024). Unsupervised ensemble-based phenotyping enhances discoverability of genes related to left-ventricular morphology. Nature Machine Intelligence, 6(3), 291–306.

Elliott, L. T., Sharp, K., Alfaro-Almagro, F., Shi, S., Miller, K. L., Douaud, G., Marchini, J., & Smith, S. M. (2018). Genome-wide association studies of brain imaging phenotypes in UK Biobank. Nature, 562(7726), 210–216.

Fan, C. C., Loughnan, R., Makowski, C., Pecheva, D., Chen, C.-H., Hagler, D. J., Jr, Thompson, W. K., Parker, N., van der Meer, D., Frei, O., Andreassen, O. A., & Dale, A. M. (2022). Multivariate genome-wide association study on tissue-sensitive diffusion metrics highlights pathways that shape the human brain. Nature Communications, 13(1), 2423.

Fu, J., Zhang, Q., Wang, J., Wang, M., Zhang, B., Zhu, W., Qiu, S., Geng, Z., Cui, G., Yu, Y., Liao, W., Zhang, H., Gao, B., Xu, X., Han, T., Yao, Z., Qin, W., Liu, F., Liang, M., … CHIMGEN Consortium. (2024). Cross-ancestry genome-wide association studies of brain imaging phenotypes. Nature Genetics, 56(6), 1110–1120.

García-Marín, L. M., Campos, A. I., Diaz-Torres, S., Rabinowitz, J. A., Ceja, Z., Mitchell, B. L., Grasby, K. L., Thorp, J. G., Agartz, I., Alhusaini, S., & Others. (2024). Genomic analysis of intracranial and subcortical brain volumes yields polygenic scores accounting for variation across ancestries. Nature Genetics, 1–12.

Glasser, M. F., Sotiropoulos, S. N., Wilson, J. A., Coalson, T. S., Fischl, B., Andersson, J. L., Xu, J., Jbabdi, S., Webster, M., Polimeni, J. R., & Others. (2013). The minimal preprocessing pipelines for the Human Connectome Project. Neuroimage, 80, 105–124.

Grasby, K. L., Jahanshad, N., Painter, J. N., Colodro-Conde, L., Bralten, J., Hibar, D. P., Lind, P. A., Pizzagalli, F., Ching, C. R. K., McMahon, M. A. B., & Others. (2020). The genetic architecture of the human cerebral cortex. Science, 367(6484), eaay6690.

Hibar, D. P., Stein, J. L., Renteria, M. E., Arias-Vasquez, A., Desrivières, S., Jahanshad, N., Toro, R., Wittfeld, K., Abramovic, L., Andersson, M., & Others. (2015). Common genetic variants influence human subcortical brain structures. Nature, 520(7546), 224–229.

Hinton, G. E., & Salakhutdinov, R. R. (2006). Reducing the dimensionality of data with neural networks. Science (New York, N.Y.), 313(5786), 504–507.

Jansen, P. R., Nagel, M., Watanabe, K., Wei, Y., Savage, J. E., de Leeuw, C. A., van den Heuvel, M. P., van der Sluis, S., & Posthuma, D. (2022). Author Correction: Genome-wide meta-analysis of brain volume identifies genomic loci and genes shared with intelligence. Nature Communications, 13(1), 3313.

Jenkinson, M., Beckmann, C. F., Behrens, T. E. J., Woolrich, M. W., & Smith, S. M. (2012). Fsl. Neuroimage, 62(2), 782–790.

Jiang, L., Zheng, Z., Qi, T., Kemper, K. E., Wray, N. R., Visscher, P. M., & Yang, J. (2019). A resource-efficient tool for mixed model association analysis of large-scale data. In bioRxiv. bioRxiv. 10.1101/598110

Kingma, D. P., Welling, M., & Others. (2019). An introduction to variational autoencoders. Foundations and Trends® in Machine Learning, 12(4), 307–392.

Littlejohns, T. J., Holliday, J., Gibson, L. M., Garratt, S., Oesingmann, N., Alfaro-Almagro, F., Bell, J. D., Boultwood, C., Collins, R., Conroy, M. C., & Others. (2020). The UK Biobank imaging enhancement of 100,000 participants: rationale, data collection, management and future directions. Nature Communications, 11(1), 2624.

Makowski, C., Wang, H., Srinivasan, A., Qi, A., Qiu, Y., van der Meer, D., Frei, O., Zou, J., Visscher, P. M., Yang, J., & Chen, C.-H. (2023). Larger cerebral cortex is genetically correlated with greater frontal area and dorsal thickness. Proceedings of the National Academy of Sciences of the United States of America, 120(11), e2214834120.

Miller, K. L., Alfaro-Almagro, F., Bangerter, N. K., Thomas, D. L., Yacoub, E., Xu, J., Bartsch, A. J., Jbabdi, S., Sotiropoulos, S. N., Andersson, J. L. R., Griffanti, L., Douaud, G., Okell, T. W., Weale, P., Dragonu, I., Garratt, S., Hudson, S., Collins, R., Jenkinson, M., … Smith, S. M. (2016). Multimodal population brain imaging in the UK Biobank prospective epidemiological study. Nature Neuroscience, 19(11), 1523–1536.

Passemard, S., Kaindl, A. M., & Verloes, A. (2013). Microcephaly. Handbook of Clinical Neurology, 111, 129–141.

Patel, K., Xie, Z., Yuan, H., Islam, S. M. S., Xie, Y., He, W., Zhang, W., Gottlieb, A., Chen, H., Giancardo, L., Knaack, A., Fletcher, E., Fornage, M., Ji, S., & Zhi, D. (2024). Unsupervised deep representation learning enables phenotype discovery for genetic association studies of brain imaging. Communications Biology, 7(1), 414.

Perlaki, G., Horvath, R., Nagy, S. A., Bogner, P., Doczi, T., Janszky, J., & Orsi, G. (2017). Comparison of accuracy between FSL’s FIRST and Freesurfer for caudate nucleus and putamen segmentation. Scientific Reports, 7(1), 2418.

Rahaman, N., Baratin, A., Arpit, D., Draxler, F., Lin, M., Hamprecht, F., Bengio, Y., & Courville, A. (2019). On the spectral bias of neural networks. International Conference on Machine Learning, 5301–5310.

Rascovsky, K., Hodges, J. R., Knopman, D., Mendez, M. F., Kramer, J. H., Neuhaus, J., van Swieten, J. C., Seelaar, H., Dopper, E. G. P., Onyike, C. U., Hillis, A. E., Josephs, K. A., Boeve, B. F., Kertesz, A., Seeley, W. W., Rankin, K. P., Johnson, J. K., Gorno-Tempini, M.-L., Rosen, H., … Miller, B. L. (2011). Sensitivity of revised diagnostic criteria for the behavioural variant of frontotemporal dementia. Brain: A Journal of Neurology, 134(Pt 9), 2456–2477.

Satizabal, C. L., Adams, H. H. H., Hibar, D. P., White, C. C., Knol, M. J., Stein, J. L., Scholz, M., Sargurupremraj, M., Jahanshad, N., Roshchupkin, G. V., & Others. (2019). Genetic architecture of subcortical brain structures in 38,851 individuals. Nature Genetics, 51(11), 1624–1636.

Shadrin, A. A., Kaufmann, T., van der Meer, D., Palmer, C. E., Makowski, C., Loughnan, R., Jernigan, T. L., Seibert, T. M., Hagler, D. J., Smeland, O. B., Motazedi, E., Chu, Y., Lin, A., Cheng, W., Hindley, G., Thompson, W. K., Fan, C. C., Holland, D., Westlye, L. T., … Dale, A. M. (2021). Vertex-wise multivariate genome-wide association study identifies 780 unique genetic loci associated with cortical morphology. NeuroImage, 244(118603), 118603.

Smith, L. N. (2018). A disciplined approach to neural network hyper-parameters: Part 1--learning rate, batch size, momentum, and weight decay. arXiv Preprint arXiv:1803. 09820.

Smith, S. M., Douaud, G., Chen, W., Hanayik, T., Alfaro-Almagro, F., Sharp, K., & Elliott, L. T. (2021). An expanded set of genome-wide association studies of brain imaging phenotypes in UK Biobank. Nature Neuroscience, 24(5), 737–745.

Strang, K. H., Golde, T. E., & Giasson, B. I. (2019). MAPT mutations, tauopathy, and mechanisms of neurodegeneration. Laboratory Investigation, 99(7), 912–928.

Subramanian, A., Tamayo, P., Mootha, V. K., Mukherjee, S., Ebert, B. L., Gillette, M. A., Paulovich, A., Pomeroy, S. L., Golub, T. R., Lander, E. S., & Mesirov, J. P. (2005). Gene set enrichment analysis: a knowledge-based approach for interpreting genome-wide expression profiles. Proceedings of the National Academy of Sciences of the United States of America, 102(43), 15545–15550.

Sudlow, C., Gallacher, J., Allen, N., Beral, V., Burton, P., Danesh, J., Downey, P., Elliott, P., Green, J., Landray, M., Liu, B., Matthews, P., Ong, G., Pell, J., Silman, A., Young, A., Sprosen, T., Peakman, T., & Collins, R. (2015). UK biobank: an open access resource for identifying the causes of a wide range of complex diseases of middle and old age. PLoS Medicine, 12(3), e1001779.

van der Meer, D., Frei, O., Kaufmann, T., Shadrin, A. A., Devor, A., Smeland, O. B., Thompson, W. K., Fan, C. C., Holland, D., Westlye, L. T., Andreassen, O. A., & Dale, A. M. (2020). Understanding the genetic determinants of the brain with MOSTest. Nature Communications, 11(1), 3512.

Vaswani, A., Shazeer, N., Parmar, N., Uszkoreit, J., Jones, L., Gomez, A. N., Kaiser, Ł., & Polosukhin, I. (2017). Attention is all you need. Advances in Neural Information Processing Systems, 30.

Vogel, J. W., Iturria-Medina, Y., Strandberg, O. T., Smith, R., Levitis, E., Evans, A. C., & Hansson, O. (2020). Spread of pathological tau proteins through communicating neurons in human Alzheimer’s disease. Nature Communications, 11(1), 2612.

Warrier, V., Stauffer, E.-M., Huang, Q. Q., Wigdor, E. M., Slob, E. A. W., Seidlitz, J., Ronan, L., Valk, S. L., Mallard, T. T., Grotzinger, A. D., Romero-Garcia, R., Baron-Cohen, S., Geschwind, D. H., Lancaster, M. A., Murray, G. K., Gandal, M. J., Alexander-Bloch, A., Won, H., Martin, H. C., … Bethlehem, R. A. I. (2023). Genetic insights into human cortical organization and development through genome-wide analyses of 2,347 neuroimaging phenotypes. Nature Genetics, 55(9), 1483–1493.

Watanabe, K., Taskesen, E., van Bochoven, A., & Posthuma, D. (2017). Functional mapping and annotation of genetic associations with FUMA. Nature Communications, 8(1), 1826.

Zhao, X., Xie, Z., Islam, Saiful, S. M., Xia, T., Chen, Cheng, & Zhi, D. (2026). TorchGWAS : GPU-accelerated GWAS for thousands of quantitative phenotypes. In arXiv [cs.DC]. arXiv. http://arxiv.org/abs/2604.21095

