## Supplementary material for "Learning dynamics of unsupervised deep learning reveal epoch-specific genetic architectures of brain morphology": Supplimentary Figures

**Supplementary Figures 1–7**

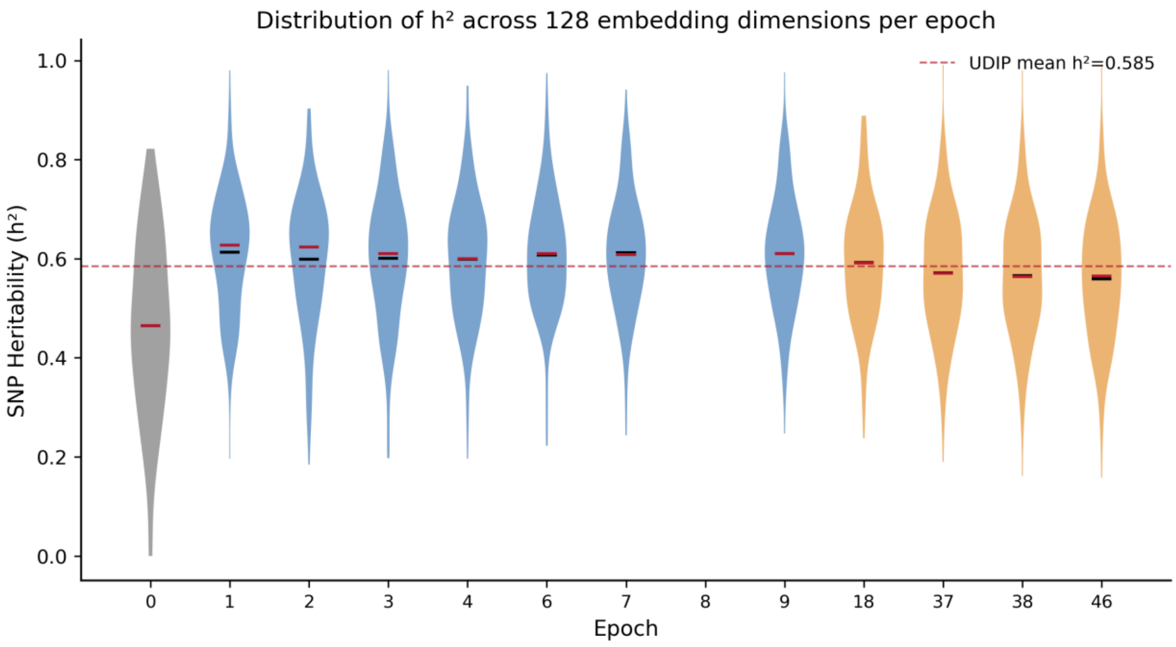


**Supplementary Figure 1 | SNP heritability distribution across informative epochs.** SNP heritability ($h^2$) across 128 embedding dimensions for each of the 13 informative epochs (GCTA --fastGWA-mlm --model-only); horizontal bar = median, dashed red line = UDIP reference mean ($h^2 = 0.585$). $h^2$ is high and stable across all epochs (grand mean 0.585), confirming that epoch-specific signals reflect genuine heritable variation. Per-epoch and per-dimension statistics in Supplementary Table 1.

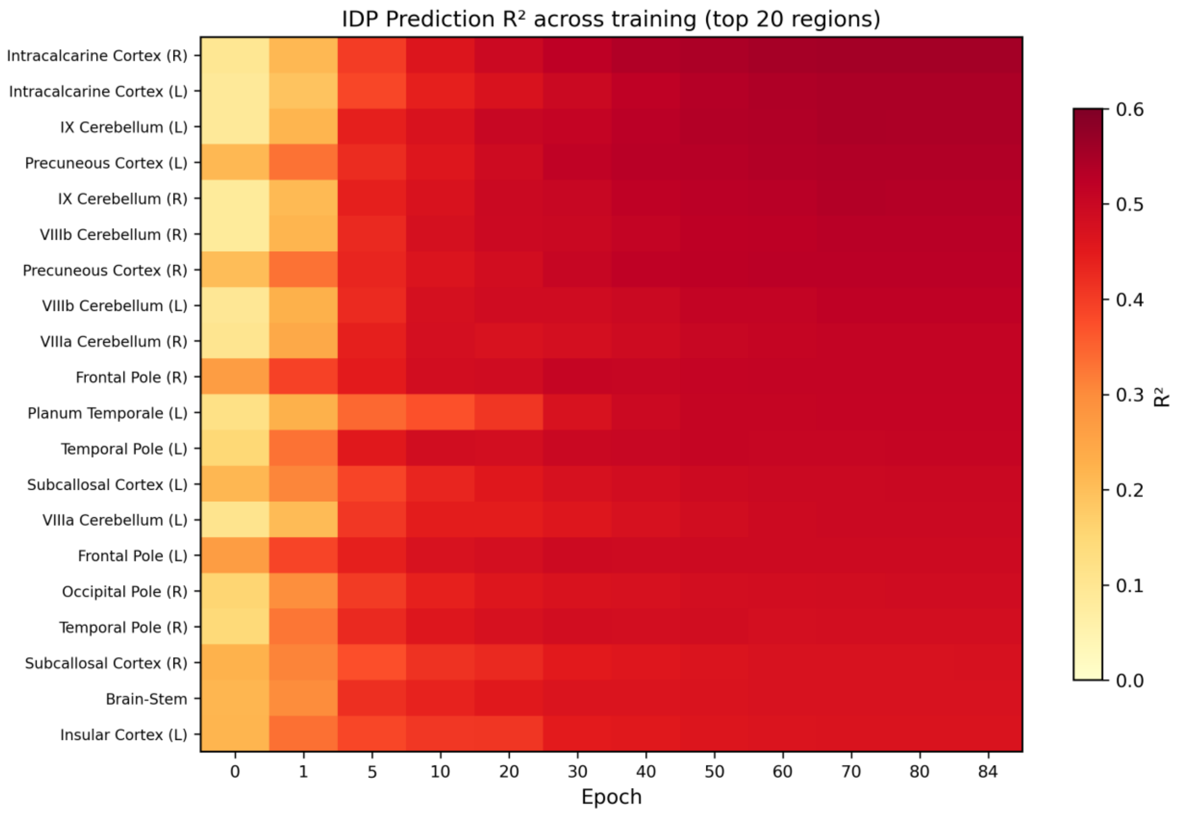


**Supplementary Figure 2 | IDP prediction $R^2$ heatmap across training epochs.** 5-fold CV OLS prediction $R^2$ for the 20 best-predicted regional grey matter volumes at 8 epochs (0, 1, 4, 9, 18, 37, 46, 84); regions sorted by peak $R^2$. Mean $R^2$ rises from 0.112 (epoch 0) to 0.352 (epoch 84); temporal lobe structures show disproportionate gains at MAPT-informative epochs (37, 46). Full $R^2$ for all 139 GM volumes in Supplementary Table 5.

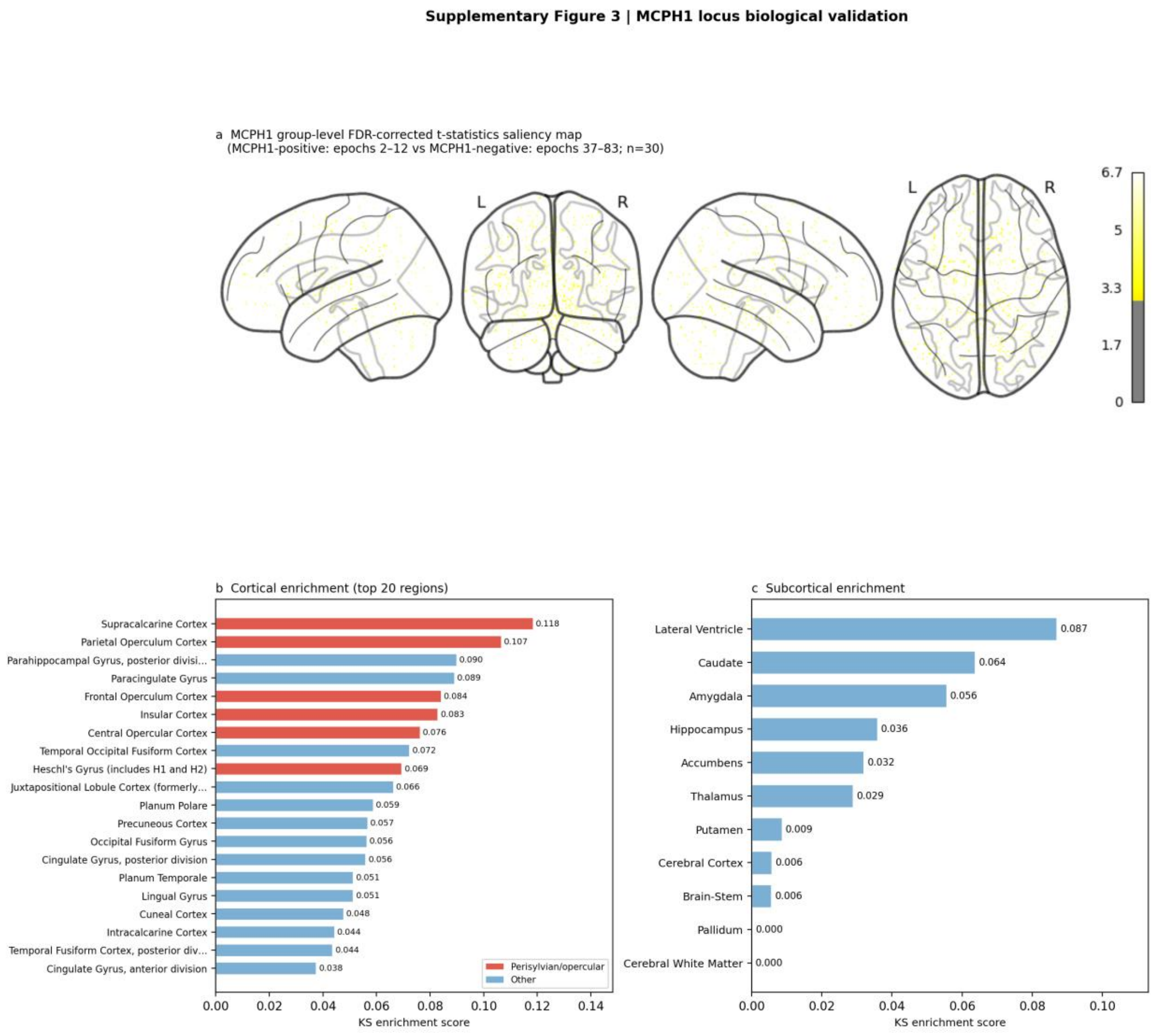


**Supplementary Figure 3 | MCPH1 locus biological validation.** MCPH1 locus (chr8p23.1, rs2979671, $P = 2.1 \times 10^{-12}$) signal is present at early epochs (2–12) and absent from epoch 13 onward. (a) Glass brain projection (left sagittal, posterior coronal, right sagittal, axial) of group-level FDR-corrected t-statistics ($n = 30$; MCPH1-positive epochs 2–12 vs MCPH1-negative epochs 37–83); 937 voxels surviving FDR $q < 0.05$ are shown ($t = 3.6$–$6.7$). (b) KS enrichment scores for 47 Harvard–Oxford cortical regions; perisylvian and opercular regions (red) dominate the top ranks — Supracalcarine Cortex (KS = 0.118), Parietal Operculum Cortex (0.107), Frontal Operculum Cortex (0.084), Insular Cortex (0.083), Central Opercular Cortex (0.076) — consistent with MCPH1's established role in perisylvian gyrification. (c) Subcortical KS enrichment (11 structures) is modest (Lateral Ventricle KS = 0.087), contrasting with the cortical signal in (b). The perisylvian pattern is spatially distinct from the temporal lobe pattern observed for MAPT (Fig. 4), confirming that early and late epochs capture distinct biological architectures. Full enrichment scores in Supplementary Table 6.

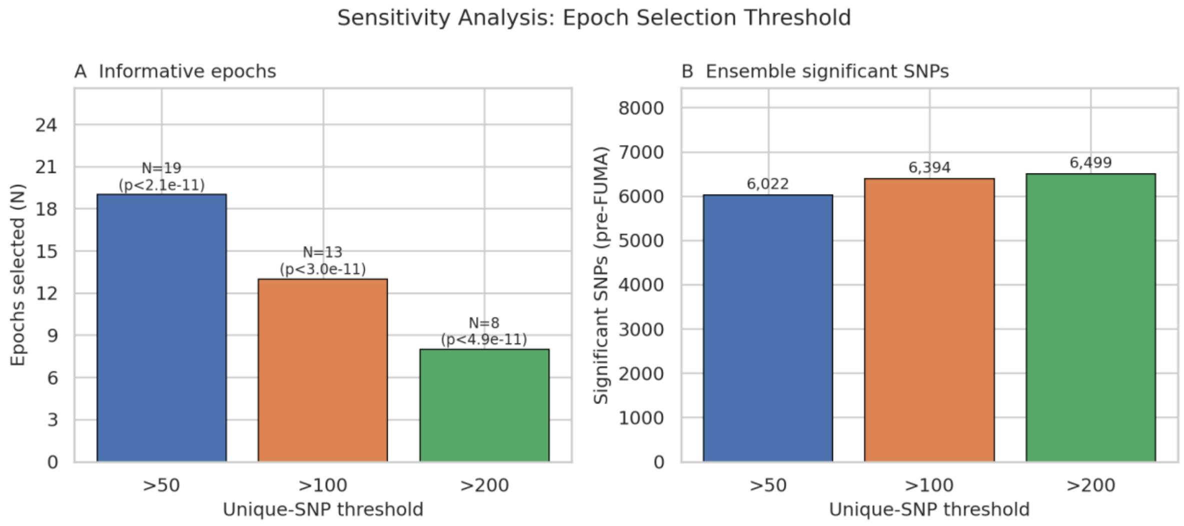


**Supplementary Figure 4 | Sensitivity analysis of the epoch selection threshold.** Genomic loci (post-FUMA clumping) under three unique-SNP thresholds for defining informative epochs: ≥ 50 SNPs (19 epochs, 86 loci), ≥ 100 SNPs (13 epochs, 90 loci; primary), ≥ 200 SNPs (8 epochs, 90 loci). The primary result (90 loci) is reproduced at threshold = 200; the slight drop to 86 loci at threshold = 50 reflects the stricter Bonferroni correction from including 19 epochs ($2.06 \times 10^{-11}$ vs $3.00 \times 10^{-11}$). Results are stable across thresholds.

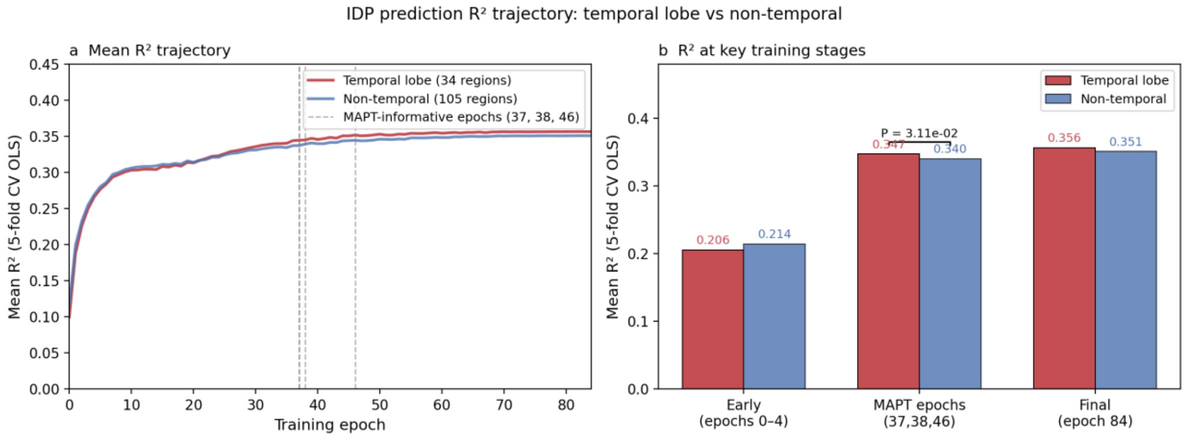


**Supplementary Figure 5 | Epoch-specific convergence of genetic and anatomical sensitivity.** (a) Mean IDP prediction $R^2$ trajectory across 85 epochs for temporal lobe (34 GM volumes; red) vs non-temporal regions (105 GM volumes; blue); dashed grey lines mark MAPT-informative epochs (37, 38, 46). (b) Mean $R^2$ at early epochs (0–4), MAPT-informative epochs (37, 38, 46), and final epoch (84). Temporal lobe gain: 0.206 → 0.347 (+69%); non-temporal: 0.214 → 0.340 (+59%); difference significant (Mann–Whitney U, $P = 0.031$). MAPT top-KS regions (Temporal Fusiform, Inferior Temporal Gyrus) reach $R^2 = 0.33$ at MAPT epochs ($\Delta = +0.13$). This cross-modal convergence at identical epochs independently validates epoch-specific genetic sensitivity.

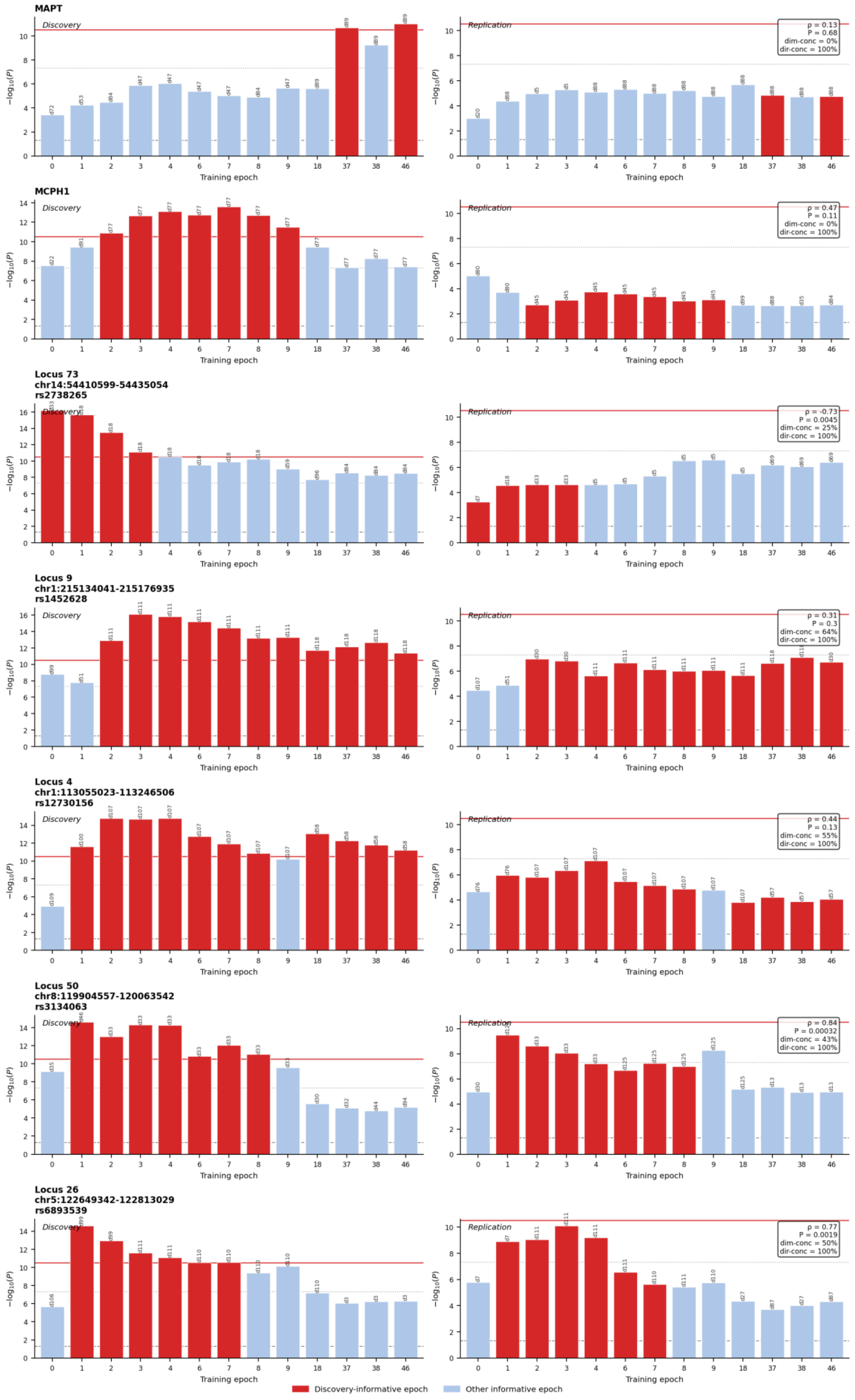

MAPT
Discovery
Replication
ρ = 0.13
P = 0.68
dim-conc = 0%
dir-conc = 100%
MCPH1
Discovery
Replication
ρ = 0.47
P = 0.11
dim-conc = 0%
dir-conc = 100%
Locus 73
chr14:54410599-54435054
rs2738265
Discovery
Replication
ρ = -0.73
P = 0.0045
dim-conc = 25%
dir-conc = 100%
Locus 9
chr1:215134041-215176935
rs1452628
Discovery
Replication
ρ = 0.31
P = 0.3
dim-conc = 64%
dir-conc = 100%
Locus 4
chr1:113055023-113246506
rs12730156
Discovery
Replication
ρ = 0.44
P = 0.13
dim-conc = 55%
dir-conc = 100%
Locus 50
chr8:119904557-120063542
rs3134063
Discovery
Replication
ρ = 0.84
P = 0.00032
dim-conc = 43%
dir-conc = 100%
Locus 26
chr5:122649342-122813029
rs6893539
Discovery
Replication
ρ = 0.77
P = 0.0019
dim-conc = 50%
dir-conc = 100%
−log10(P)
Training epoch
Discovery-informative epoch
Other informative epoch

**Supplementary Figure 6 | Epoch-specificity replication in an independent cohort.** Per-epoch $-\log_{10}(P)$ profiles for MAPT, MCPH1, and the five top-ranked TRACE-unique loci, shown in discovery (left; $N \approx 23k$) and replication (right; $N = 12{,}258$). Red bars indicate discovery-informative epochs (ensemble Bonferroni $p < 3 \times 10^{-11}$; red horizontal line). In all seven loci, the lead SNP is nominally significant ($p < 0.05$) at 100% of its discovery-informative epochs in replication, with 100% effect direction concordance. Spearman $\rho$ (epoch profile concordance between discovery and replication) and direction concordance are annotated on each replication panel. Loci are identified by FUMA locus ID, genomic coordinates, and lead SNP. Full per-locus results in Supplementary Tables 8–9.

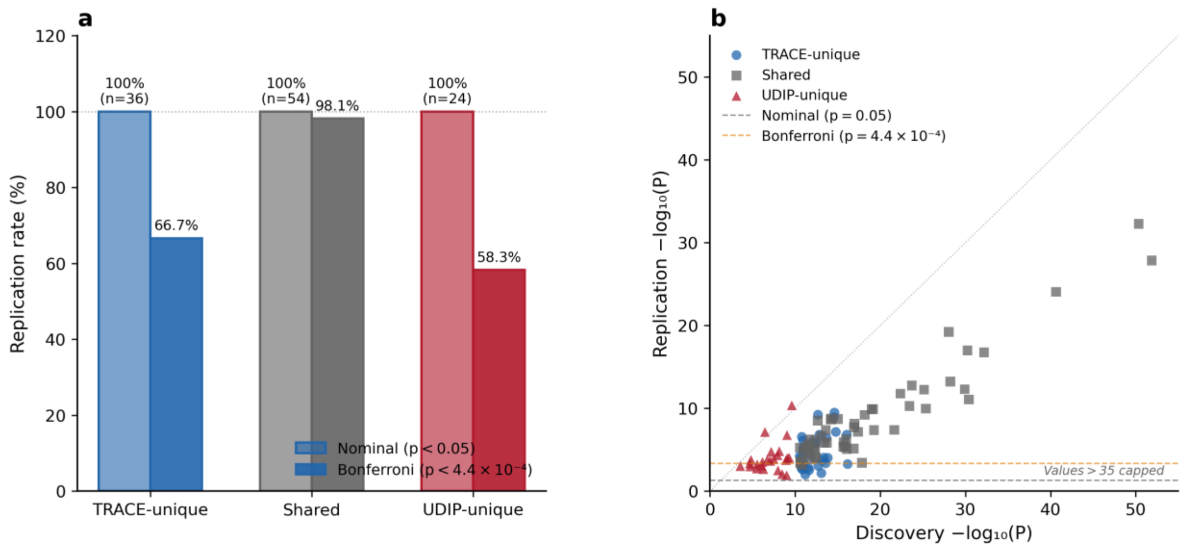


**Supplementary Figure 7 | Replication of all 114 discovered loci.** (a) Nominal and Bonferroni replication rates by locus category (TRACE-unique, Shared, UDIP-unique). All 114 loci replicated nominally (100%); shared loci replicated at the highest Bonferroni rate (98.1%), reflecting detection by two independent methods. (b) Scatter plot of discovery vs replication $-\log_{10}(P)$ for all 114 loci, colored by category (TRACE-unique: blue circles; Shared: grey squares; UDIP-unique: red triangles). Dashed lines indicate nominal ($p = 0.05$) and Bonferroni ($p = 4.4 \times 10^{-4}$) thresholds. Stronger discovery signal predicts stronger replication signal, consistent with a power explanation for the lower Bonferroni rate among method-specific loci. Three discovery values capped at 35 for display. Per-locus results in Supplementary Table 8.
